# Low but genetically variable male remating ability in a tropical *Drosophila* despite substantial fitness benefits

**DOI:** 10.1101/504035

**Authors:** Andrew D. Saxon, Natalie E. Jones, Eleanor K. O’Brien, Jon R. Bridle

**Affiliations:** School of Biological Sciences, Life Sciences Building, University of Bristol, Bristol. BS8 1TQ. U.K

**Keywords:** Male remating traits, reproductive success, sperm allocation, heritability, offspring quality

## Abstract

Mating success is the main source of fitness variation in males, meaning that males should capitalise on all opportunities for mating. Strong selection on male mating success should also reduce genetic variation in male mating traits relative to other traits. We quantified mating latency, mating duration and productivity in males of the tropical fruitfly, *Drosophila birchii*, from 30 isofemale lines collected from across two elevational gradients, when they were given opportunities to mate with up to four females consecutively. Male remating rates were low compared to other *Drosophila* (only 14 – 27% of males achieved a fourth mating), with mean mating durations approximately doubling across successive copulations. However, although successive remating produced progressively fewer offspring, it consistently increased overall male reproductive success, with males that mated four times more than doubling offspring number compared to males mating only once. We also found no reduction in the productivity of sons emerging from later matings, indicating a sustained cumulative fitness benefit to remating. Heritable variation was observed for most traits (*H*^2^ = 0.035 – 0.292) except mating latency, but there was no divergence in trait means with elevation. The observed restricted remating ability of male *D. birchii*, despite the clear benefits of remating, may be due to a low encounter rate with females in the field, leading to high investment per gamete (or ejaculate). However, it remains unclear why genetic variation in these traits is high, given we observe no variation in these traits across elevational gradients known to affect local population density.

## 1. Introduction

Male sexual behaviour is determined by a trade-off between current and future mating success (Parker, 1982). Males should invest more in initial copulations if female mating frequency and/or encounter rate is low (Reinhold *et al*, 2002). However, if sperm is costly and male mating rate is high, strategic allocation of resources over successive matings (and successive females) is expected (Byrne and Rice, 2006; Engqvist and Sauer, 2001). It has been observed that males from several taxa partition sperm across consecutive females (Birkhead, 1991; Perez-Staples and Aluja, 2006; Pitnick and Markow, 1994; Rondeau and Sainte-Marie, 2001; Shapiro and Giraldeau, 1996; Svard and Wiklund, 1986). Understanding the evolution of male mating traits therefore demands measurement of the size of fitness benefits that males gain from multiple matings compared to their first mating, and how this varies among males and their sons (Wedell and Tregenza, 1999). Such variation in male life history within and among species determines reproductive skew, sexual conflict and dimorphism, all of which have profound implications for the ecology and evolution of populations.

Sexual activity reduces male lifespan (Partridge and Farquhar, 1981; Van Voorhies, 1992) and incurs significant energy, time and risk-related costs (Bretman *et al*, 2013b; Dowling and Simmons, 2012; Hayward and Gillooly, 2011). These include the cost of searching and competing for females (Dewsbury, 1982), perceiving female signals (Harvanek *et al*, 2017), complex courtship displays (Connolly *et al*, 1969; Cordts and Partridge, 1996) and prolonged copulation duration or extended mate guarding to maximise paternity share (Bretman *et al*, 2013a; Mazzi *et al*, 2009). Sperm production can also be energetically expensive (Dewsbury, 1982; Hayward and Gillooly, 2011; Paukku and Kotiaho, 2005), especially in species that generate large sperm (Pitnick, 1996). Sperm depletion may therefore limit male reproductive success if mating opportunities are frequent (Hines *et al*, 2003; Pitnick and Markow, 1994; Preston *et al*, 2001; Wedell *et al*, 2002). This means that if opportunities for remating are common in nature, rapid ejaculate replenishment should be under strong directional selection (Trivers, 1972).

Although larger and more numerous ejaculates generally increase fitness (Simmons, 2001), sperm quality affects male fitness independently of sperm quantity (Pattarini *et al*, 2006; Snook, 2005). Declines in sperm quality (e.g. motility, viability) occur over rapid sequential ejaculations in rabbits (*Oryctolagus cuniculus*), especially as ejaculates become depleted (Ambriz *et al*, 2002). Variation in sperm quality among males also results in differences in the quality of offspring (Alavioon *et al*, 2017; Hosken *et al*, 2003; Siva-Jothy, 2000). For example, in male guppies (*Poecilia reticulata*), declines in sperm competitive ability are associated with reduced reproductive success of sons (Gasparini *et al*, 2017). Offspring from later matings may therefore have reduced fitness compared to those from initial matings.

In *Drosophila*, sperm are transferred in a complex seminal fluid comprising many costly components (Dewsbury, 1982; Perry *et al*, 2013), including accessory gland proteins (Acps) that can affect female reproductive behaviour (Gillott, 2003). However, producing these seminal components may also limit the number and size of ejaculates males can produce (Lefevre and Jonsson, 1962; Linklater *et al*, 2007; Reinhardt *et al*, 2011; Sirot *et al*, 2009; Wigby *et al*, 2009), and the replenishment of seminal fluids (or maturation of sperm) is associated with delays before male remating in other insects (Vahed, 2007). Males also adjust copulation duration and ejaculate quantity in the presence of rival males, presumably associated with the likelihood of sperm competition (Gage and Baker, 1991). Edvardsson and Canal (2006) suggest that declines in copulation duration are associated with decreasing ejaculate transfer, with reductions in mating duration observed in males of several *Drosophila* species over consecutive matings (Linklater *et al*, 2007; Singh and Singh, 2013; Singh and Singh, 2000).

Selection to prioritise current versus future mating may also vary across environments, and with population density. In *Drosophila*, variation in ejaculate size correlates with variation in female fecundity, mating status and age (Lupold *et al*, 2011; Wedell *et al*, 2002), all of which may vary across environments (Chapman and Partridge, 1996). Ecological conditions also determine mating patterns (Gromko and Markow, 1993), due to trade-offs between stress tolerance and reproduction (Marshall and Sinclair, 2010). For example, low resource availability may reduce male mating rates (Blay and Yuval, 1997) and their ability to produce sperm (Gage and Cook, 1994). Latitudinal clines in male mating traits indicate that environmental variation can determine such allocation patterns (Chahal *et al*, 2013).

Although selection on male mating traits is likely to vary across environments, mating success traits should have lower heritabilities than morphological or physiological traits because of their close association with fitness. Alternatively, low heritability estimates in mating traits could result from their higher sensitivity to environmental variation (Price and Schluter, 1991). Understanding genetic variation in mating traits is key to predicting the ecological factors that determine their evolution across species’ ranges, and therefore the range of environmental conditions where their populations can persist (Nadeau *et al*, 2017).

Studies of male mating traits in *Drosophila* mainly focus on cosmopolitan species that are easy to rear in the laboratory and occur at high density throughout their ecological range (Gowaty, 2012). Tropical species are less well investigated despite typically having narrower abiotic and biotic tolerances which should have direct and indirect effects on reproduction (Markow and O’Grady, 2008), and so provide important information on how ecological and demographic factors affect he evolution of mating behaviour. In this study, we assayed the mating behaviour and reproductive success of males of the tropical fly *Drosophila birchii*, to explore the effects of its previously observed low male remating rate (when used in paternal half-sib breeding designs), compared to its close relatives *Drosophila serrata* (Sgro and Blows, 2003) and *D. bunnanda* (Van Homrigh *et al*, 2007). This species is limited to tropical rainforest in north-eastern Australia and New Guinea and has an elevational range from ∼0 – 1100 metres, which is characterised by considerable variation in population density across elevation (Bridle *et al*, 2009; O’Brien *et al*, 2017). It also displays complex male courtship behaviour (Hoikkala *et al*, 2000) and substantial reductions in male mating success under fluctuating versus constant thermal regimes (Saxon *et al*, 2018).

We assessed genetic variation across the elevational range of this species (at two locations in northern Queensland), for the following mating and reproductive traits: (*i*) latency to achieve copulation, (*ii*) duration of copulation, (*iii*) number of offspring produced with each successive copulation, and (*iv*) total number of offspring sired, when initially virgin males were presented with four virgin females sequentially. We then tested how this variation in male mating traits affects male reproductive success, both in terms of offspring and grand-offspring. Surprisingly, the remating ability of *D. birchii* was found to be remarkably low across all habitats, despite higher remating rates consistently increasing numbers of offspring and grand-offspring and significant levels of genetic variation in key male mating traits.

## 2. Materials and Methods

### (a) Establishment of isofemale lines

*Drosophila birchii* isofemale lines (called ‘lines’ hereafter) were founded from field-mated females collected from two low or high elevation sites along two gradients (eight sites in total): Mount Edith (Elevation: ∼600 – 1100 m, 17°6’S, 145°38’E) and Mount Lewis (Elevation: ∼20 – 900 m, 16°35′S 145°17′E), in Queensland, Australia in 2011. Flies were collected using banana baited buckets, sampled daily using fine sweep nets and sorted under a microscope using light CO_2_ anaesthesia. Females were placed individually in vials to lay, for 5 – 10 days. Each line was maintained across 3 – 4 40 ml vials containing 10 ml of *Drosophila* potato food medium (agar, instant mashed potato powder, raw sugar, inactive yeast, propionic acid, nipagin supplemented with live yeast) at ∼100 individuals per generation for each line. A ‘mass-bred’ stock was established by combining 10 male and 10 female flies from each line, as a genetically mixed background population to provide a source for test females to assess the mating behaviour and fitness of focal males. Mass-bred stocks were reared in 400 ml bottles with 100 ml of *Drosophila* medium and mixed between generations. All lines and the mass-bred population were maintained with non-overlapping generations at 19 °C on a 12:12-h light:dark cycle at 60% relative humidity prior to the experiments. Mount Lewis experiments were conducted after ∼25 laboratory generations, Mount Edith ∼50 generations.

### (b) Experiment I: Assaying male remating and productivity

Ten lines were used for the Mount Edith assay, with five lines originating from two high elevation sites and five from two lower elevation sites. Twenty lines were used for the Mount Lewis assay, with 10 originating from two high and two low sites. Prior to the experiment, line stocks and the mass-bred population were reared at a constant 25 °C, 12:12 hour light:dark cycle for two generations to randomise and minimise any transgenerational effects attributable to maternal condition. The experimental males and background females were reared at minimal density conditions to minimise larval competition. Five male and five female flies mated and laid eggs in 10 ml of standard *Drosophila* medium for three days. Parental flies were then removed and pupation card (75 × 30mm) inserted into the vial.

On eclosion, flies were anaesthetised under CO_2_ and sexed using a Leica (MZ9.5) microscope. 30 males for each of 10 lines (*N* = 300) from Mount Edith and 20 males from 20 lines (*N*= 400) from Mount Lewis were collected, along with mass-bred females, within 12 hours of emergence to ensure virgin status. All flies were held in single-sex vials for six days, at 25 °C in low density vials (maximum of 10 flies per vial) with fly food medium *ad libitum*, before the mating assay commenced.

#### Mating assay

The assay was conducted at a constant 25 °C and began within an hour of the daylight period, to coincide with time of peak activity in *Drosophila*. Each male was placed in a vial with 8 ml of standard fly medium. A virgin female was placed with the focal male and the start time noted. If mating was initiated, the time was recorded to give time to copulation (latency), as was the end mating time (duration). Following copulation, the male was moved to a fresh vial and presented with a new female. This process continued with up to four matings allowed for each male, with males given up to two hours to mate with each female. If no mating occurred, then the male was recorded as ‘not mated’ and the assay concluded for that male. The assay took place over five days with 60 (for Mount Edith) and 80 (for Mount Lewis) male flies assayed per day, using equal numbers from each line. The order of males was randomised within each block to ensure that there was no systematic bias among lines due to diurnal effects. Blind ID codes were used to avoid observer bias.

Mated females were left in the vials for five days, which ensured that they had laid all fertilised eggs, given that the lack of spermathecae in *D. birchii* females means that sperm storage is minimal (R. Snook, personal communication, 2014). The female was then removed and a pupation card added. At least 20 days after mating, the total productivity (number of offspring) of each successful mating for each male was recorded.

### (c) Experiment II: Assaying the fitness of sons derived from successive paternal matings

Ten virgin males (sons) were collected from each of the first to fourth (1 – 4) paternal matings of focal males (sires) in Experiment I that had achieved the maximum four matings. Sons were derived from seven of the ten Mount Edith isofemale lines (*N* = ∼280). Each six-day old male was placed in a single vial of 8 ml of fly medium with a single, virgin mass-bred female of the same age (± 12 hours). The pair were left for three days to mate under the same control conditions as Experiment I. The male was then removed and the female was left to lay for a further five days. The female was then removed and pupation card inserted. After all offspring had emerged, the mean productivity of matings from these sons was assayed with respect to the order of mating and their paternal line.

### (d) Statistical analysis

The R (R Core Team, 2016) package *lme4* was used to fit linear mixed models to test for effects of successive matings on male mating traits (latency to mate, duration of mating, number of offspring per mating, total offspring per male). Separate models were fitted for each trait and gradient. All data were untransformed because trait data were normally distributed. Mating number and elevation of origin of line (high or low) were included as fixed factors and focal male nested within isofemale line were specified as random effects. For total offspring per male, only elevation of origin of line was used as a fixed factor. The significance of each factor in the model was determined using likelihood ratio tests to compare the full model with a model where that factor had been removed (Experiment I). *P*-values were corrected for multiple comparisons following the False Discovery Rate method of Benjamini and Hochberg (1995), allowing an FDR of 0.05. Linear mixed models were also run to analyse overall differences in the mean proportion of males per line that attained each successive mating and the mean cumulative number of offspring per line at each mating. Mating number was used as a fixed factor with line as a random effect. For Experiment II, a linear mixed model with mating number as a fixed factor and line as a random effect was used to analyse the productivity of sons derived from successive paternal mating events when mating with a single, virgin female from the mass-bred stock.

To estimate broad-sense heritability (*H*^2^) for mating traits, between line variances from the linear mixed models were used (Experiment I), using within (*V*_w_) and between (*V*_b_) line variance components derived from models fitted using REML. This was an overall estimate for each gradient as the small number of lines per elevation precluded estimates for high and low elevation separately. Inbreeding coefficients (*F*_t_) (with the assumption of full-sibship) for the number of generations and population size (∼100) at which lines had been held at in the laboratory were calculated using the method of Falconer and Mackay (1996). *H*^2^ was then calculated following Hoffmann and Parsons (1988)(see SM). The significance of the *H*^*2*^ was evaluated from the significance of the between-line variance component in the linear mixed models.

## 3. Results

### (a) Experiment I: Variation in male mating traits over successive matings

In Experiment I, 76% of males at Mount Edith attained a first mating (*N* = 220 males), with only 27% reaching a fourth. 19% only mated once. 63% of males at Mount Lewis attained a first mating (*N* = 166 males), while only 14% attained a fourth mating, with 14% of males mating only once (Figure 2). Mean cumulative productivity, while significantly increasing across successive matings, slowed its rate of increase with each copulation (Fig. 3). For Mount Edith lines, mean productivity per mating (for all males) showed a 64% decrease from mating 1 to mating 4. Focal males that mated only once had mean total offspring of 43.02 (standard error ± 5.176), while males that mated four times had a mean total cumulative productivity of 145.07 (± 18.305). For Mount Lewis lines, mean productivity per mating declined by 72% from mating 1 to mating 4. Males that mated only once had mean total offspring of 63.97 (± 6.010), while males reaching a fourth mating had a mean total productivity of 133.31 (± 7.041). The subset of males that reached a fourth mating showed a similar pattern of declining productivity to the pattern observed for all males (Figure SM1).

**Figure 1.**
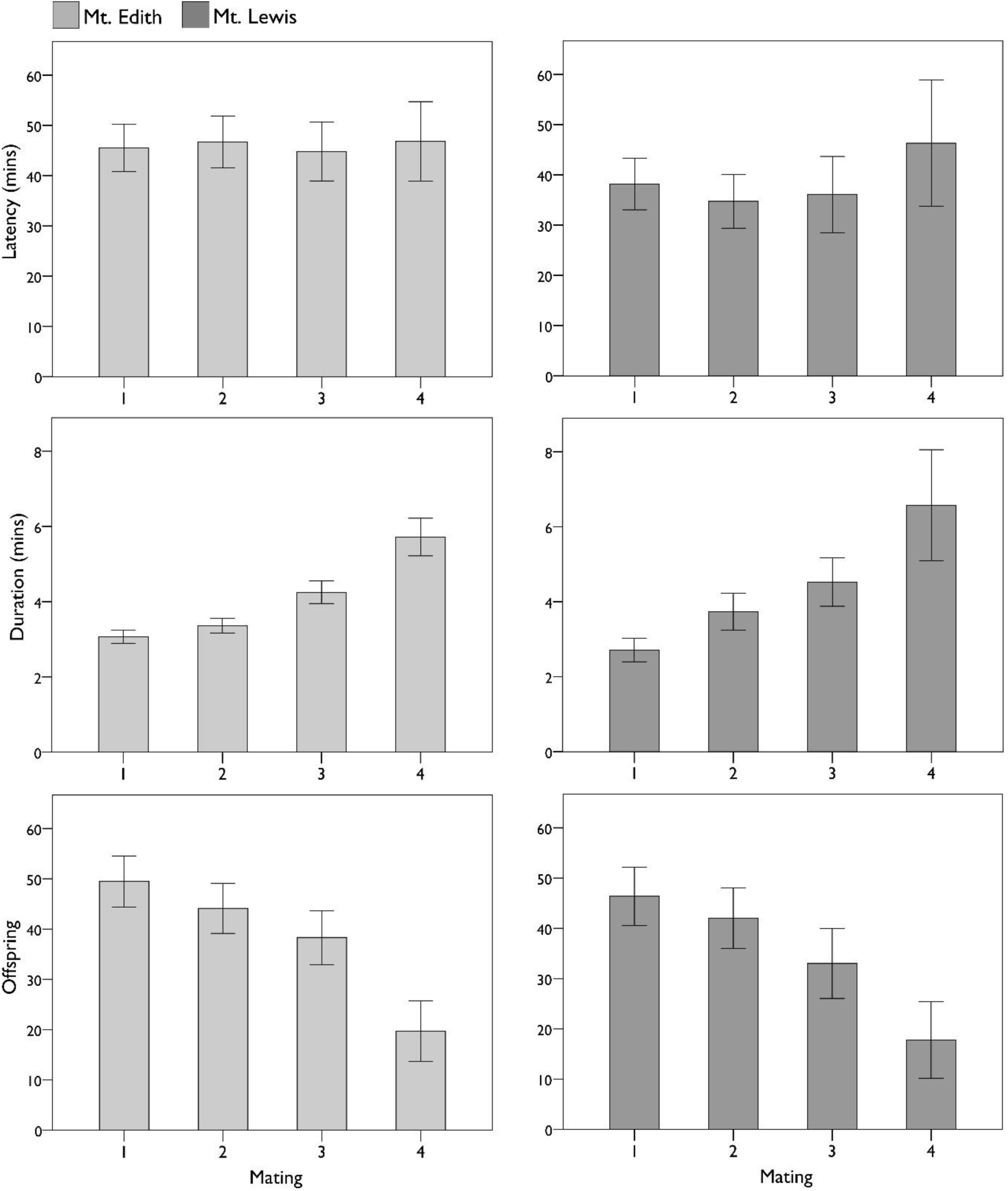
Increased mating duration and reduced productivity with successive matings: Mean mating latency, mean mating duration and mean number of offspring (productivity) for all isofemale lines for each successive male mating (1 – 4). Data from Mount Edith lines are shown in light grey (left), Mount Lewis lines shown in dark grey (right). Error bars indicate 95% confidence intervals.

**Figure 2.**
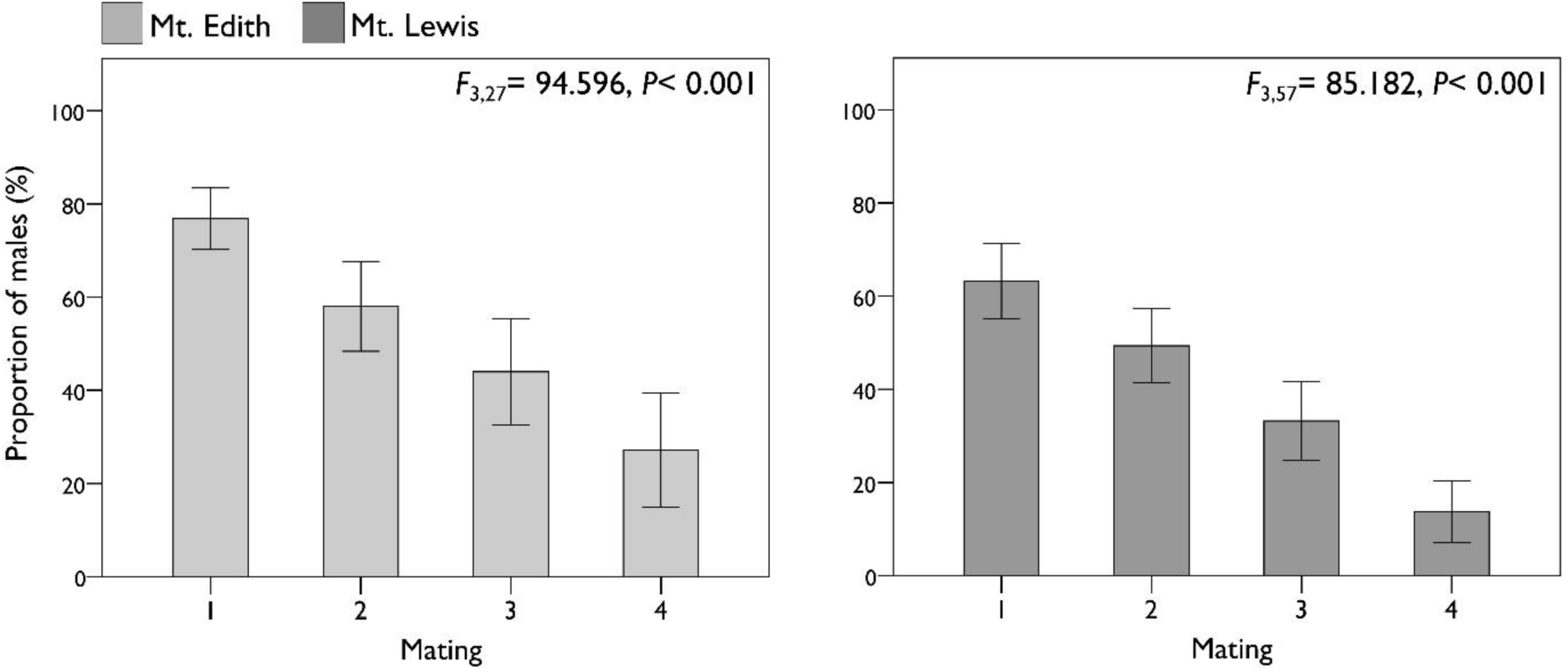
Mean proportion (%) of males per line reaching each mating. Mount Edith (left) and Mount Lewis (right) with 95% confidence intervals.

**Figure 3.**
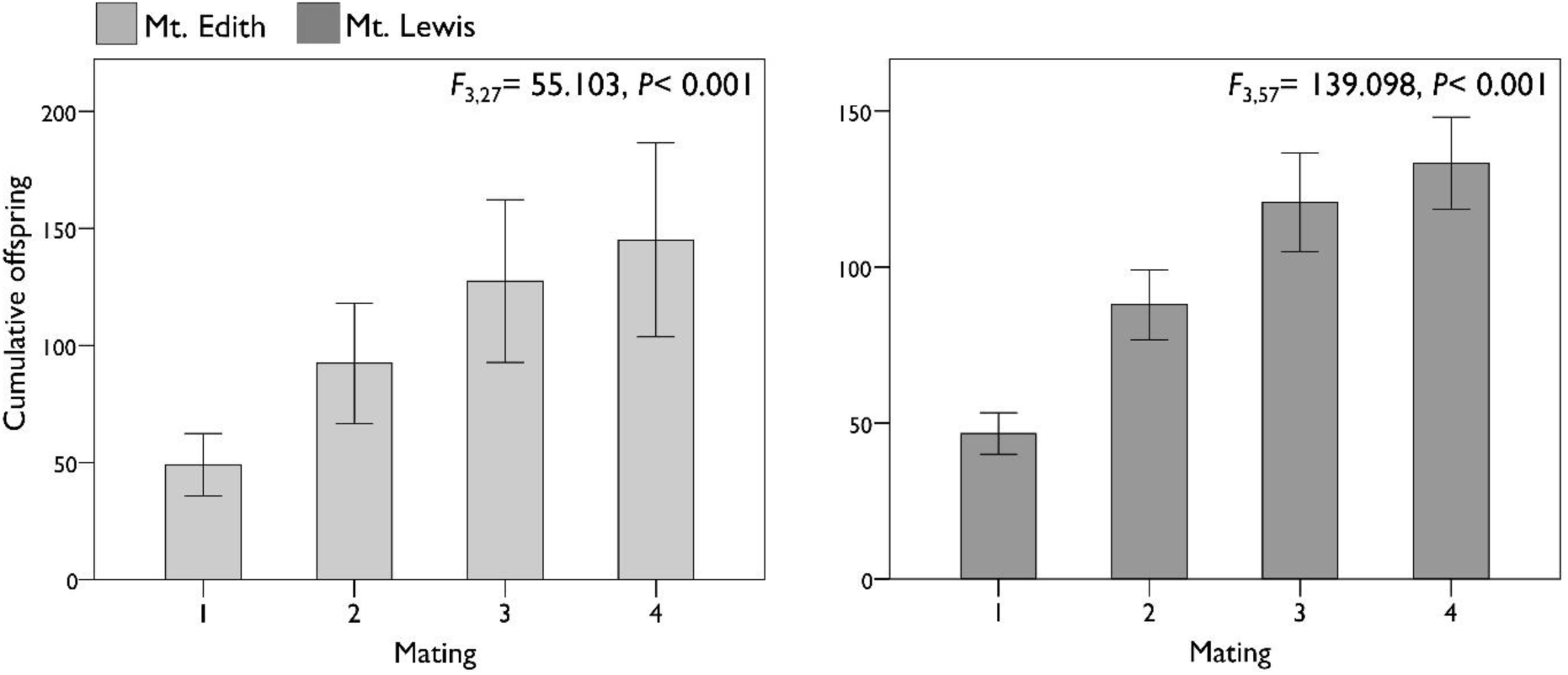
Cumulative number of offspring produced per line with successive matings. Mount Edith (left) and Mount Lewis (right) with 95% confidence intervals.

Latency to mating did not vary with successive matings for line from either gradient. Between-line (genetic) variance in this trait was also not observed at either gradient. However, both mating duration and productivity showed a highly significant change across successive matings for both gradients. Mating duration increased approximately twofold and productivity more than halved over the four successive matings (Figure 1; Table 1). Line also showed significant variance in these traits (with the exception of mating duration in the Mount Lewis lines), but there was no detectable difference between lines originating from high and low elevation sites for any of these traits. Broad-sense heritabilities (*H*^2^) for variance components in the Mount Edith and Mount Lewis (Table 2) lines gave low to moderate estimates. Mating duration gave low *H*^2^ values (0.035 – 0.126), although the estimate did not significantly differ from zero at Mount Lewis. *H*^2^ estimates were significantly different from zero for number of offspring produced per mating, ranging from 0.069 – 0.229. Total productivity per male exhibited significant heritability at Mount Edith (0.292), but not at Mount Lewis.

**Table 1.** Linear mixed effects analyses for male mating traits in Mount Edith and Mount Lewis lines. Mating (1 – 4) and elevation of origin of isofemale line are fixed effects, with nested random effects of isofemale line and focal male. *P*-values were obtained by likelihood ratio tests of the full model with the effect against the model with the effect excluded (*ns*= not significant, *adjusted *P*≤ 0.05, \*\**P* ≤ 0.01, \*\*\**P* ≤ 0.001).

| Mating trait | Gradient | Fixed effect |  |  |  | Random effects |  |  |
| --- | --- | --- | --- | --- | --- | --- | --- | --- |
| | | Predictor | $\chi^2$ | df | P | Variance component | Variance | P |
| Latency | Mt. Edith | Mating | 0.5111 | 3 | ns | Isofemale line | 28.61 | ns |
| | | | | | | Male | $9.803 \times 10^{-13}$ | ns |
|  |  | Elevation | 2.097 | 1 | ns | Residual | 1162 | - |
|  | Mt. Lewis | Mating | 4.0038 | 3 | ns | Isofemale line | 10.15 | ns |
|  |  |  |  |  |  | Male | 39.95 | ns |
|  |  | Elevation | 0.7601 | 1 | ns | Residual | 1031.17 | - |
| Duration | Mt. Edith | Mating | 184.84 | 3 | *** | Isofemale line | 0.08798 | * |
|  |  |  |  |  |  | Male | 0.04592 | ns |
|  |  | Elevation | 0.653 | 1 | ns | Residual | 3.00321 | - |
|  | Mt. Lewis | Mating | 64.142 | 3 | *** | Isofemale line | 0.7751 | ns |
| | | | | | | Male | $6.507 \times 10^{-14}$ | ns |
|  |  | Elevation | 0.5598 | 1 | ns | Residual | 8.535 | - |
| Offspring per mating | Mt. Edith | Mating | 76.697 | 3 | *** | Isofemale line | 221.5 | *** |
|  |  |  |  |  |  | Male | 106.3 | ns |
|  |  | Elevation | 1.9363 | 1 | ns | Residual | 958.3 | - |
|  | Mt. Lewis | Mating | 18.959 | 3 | *** | Isofemale line | 48.47 | * |
|  |  |  |  |  |  | Male | 184.27 | * |
|  |  | Elevation | 0.051 | 1 | ns | Residual | 1012.86 | - |
| Total Offspring | Mt. Edith | Elevation | 1.8045 | 1 | ns | Isofemale Line | 1831 | *** |
|  |  |  |  |  |  | Residual | 5809 | - |
|  | Mt. Lewis | Elevation | 0.094 | 1 | ns | Isofemale Line | 365.1 | ns |
|  |  |  |  |  |  | Residual | 3148.5 | - |

**Table 2.**
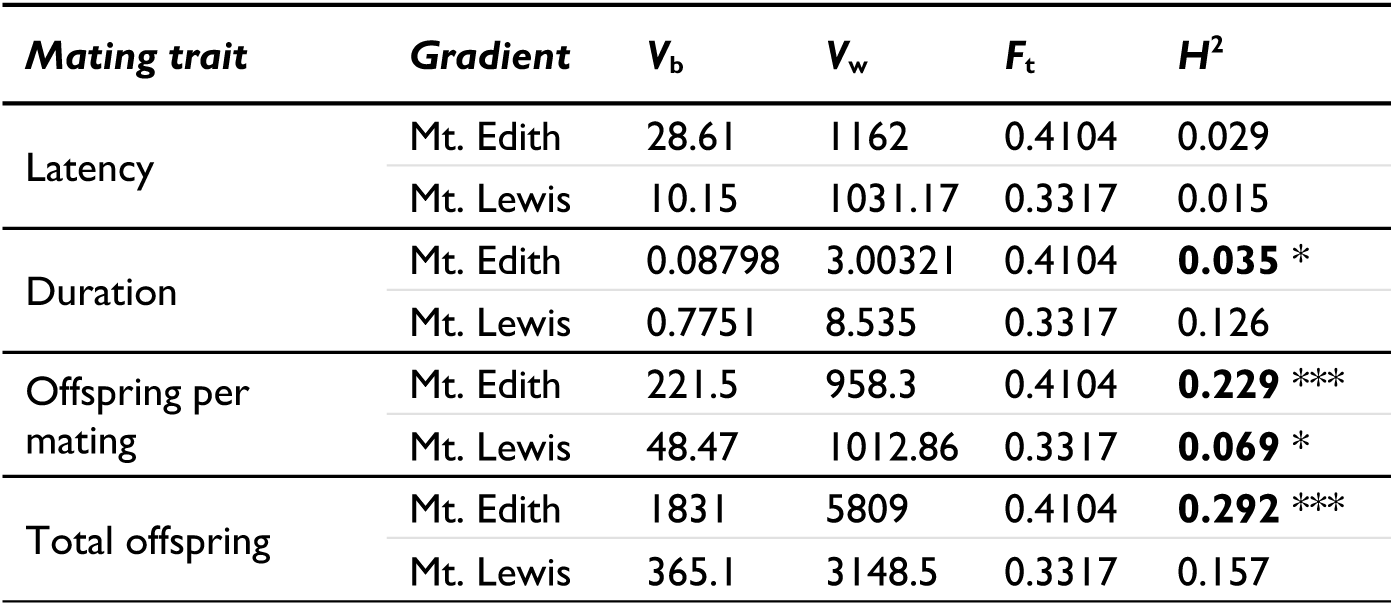
Broad-sense heritability (*H*^2^) estimates for mating traits for Mount Edith and Mount Lewis lines. Calculated from between (*V*_b_) and within (*V*_w_) line variance components and inbreeding coefficient (*F*_t_) for number of laboratory generations. Significant *H*^2^ estimates are in **bold.**

### (b) Experiment II: Fitness of sons from successive paternal matings

There were no significant differences in the productivity of sons derived from successive paternal matings in Experiment II (Fig. 4; *χ*^2^ = 0.2675, df = 1, *P* = *ns*). There was also no significant variation in productivity among sons attributable to isofemale line used (Variance = 6.65 × 10^-16^, *P* = *ns*; Residual = 537.1). Mean number of offspring produced from mating with a single, virgin female was similar for sons obtained from sire matings 1 to 4: Mating 1 = 73.3 (Standard Error ± 2.59), Mating 2 = 75.4 (± 2.96), Mating 3 = 73.5 (± 3.24), Mating 4 = 76.2 (± 2.98).

**Figure 4.**
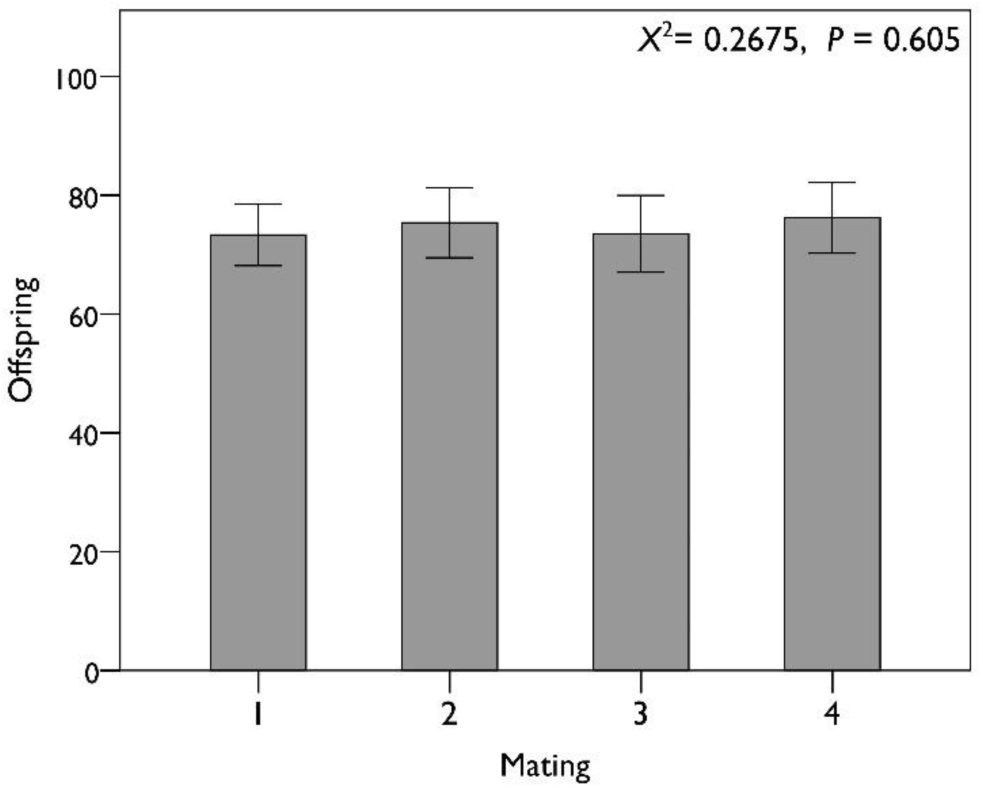
No decline in reproductive success of male offspring from later matings. Number of (grand) offspring produced by sons derived from successive paternal matings (1 – 4) when mating with a single, virgin female (with 95% confidence intervals) for Mount Edith lines.

## 4. Discussion

The substantial variation in male mating strategies observed among *Drosophila* species (Gowaty *et al*, 2003; Singh and Singh, 2013) is likely to correlate with ecological parameters that relate to the risk of sperm competition, likelihood of repeated mating opportunities, and the energetic cost of mating relative to resistance to factors such as temperature stress and pathogen or parasitoid exposure. Here, we demonstrate that males of the rainforest specialist *Drosophila birchii* exhibit a much lower remating rate than cosmopolitan species for which data are available. For example, *D. melanogaster* males will copulate with 10 females per day given the opportunity (Gromko, 1992; Mossige, 1955), while *D. hydei* can mate up to 10 times in two hours (Markow, 1985). By contrast, only 27% and 14% of *D. birchii* males, from Mount Edith and Mount Lewis lines respectively, mated a fourth time within the single day of our experiments, with only 58% and 49% mating more than once (Fig. 2). This was despite the two to threefold increase in the total number of offspring produced by males that mated four times compared to those that mated only once (Fig. 3). In addition, unlike many *Drosophila* species, female *D. birchii* have no spermathecae and therefore cannot store sperm (R. Snook, personal communication, 2014). There is also strong last male precedence in paternity for *D. birchii* when females remate (E.K. O’Brien, unpublished data, 2017). Both of these, coupled with the sustained increases in productivity where remating occurs, should make remating consistently beneficial to male fitness, even when other males are around. These observations therefore make this observation of low male remating rates especially surprising.

*Drosophila birchii* are often found at low densities in their rainforest habitat (Bridle *et al*, 2009; O’Brien *et al*, 2017), compared to other tropical drosophilids. This means conspecific interactions (especially the female encounter rate for males) may be relatively rare, which is likely to shape resource allocation across matings (Aspi and Hoffmann, 1998; Shelly and Bailey, 1992; Willis *et al*, 2011). In particular, reduced risk of sperm competition may favour reductions in sperm quantity (Bjork *et al*, 2007), in favour of increased investment per gamete (Snook, 2005) and reduced allocation across multiple matings (Ingleby *et al*, 2010). Reductions in production of seminal fluid proteins also occur in less sexually competitive environments (Wigby *et al*, 2016).

Ejaculate components such as sperm (Wedell *et al*, 2002) or seminal fluids (Wigby *et al*, 2009) can limit male remating. Spermiogenesis in male *Drosophila melanogaster* takes five days (at 25 °C) (Fabian and Brill, 2012) and mated males with depleted seminal fluid proteins require three days of sexual inactivity before transferring equivalent quantities again (Sirot *et al*, 2009). Although sperm size in *D. birchii* is not known, long flagella and long ventral receptacles in females (R. Snook, personal communication, 2014) suggest that male investment in sperm is high (Markow, 2015). If energetic costs of mating are high for male *D. birchii*, successive mating events may increase male choosiness, as the relative cost of mating rises with male resource depletion (Byrne and Rice, 2006). In this study however, no variation was found in latency to copulation across matings (Fig. 1), suggesting no increase in male choosiness (i.e. inclination to remate) (Engqvist and Sauer, 2001) or reduction in male attractiveness to females (Taylor *et al*, 2007).

Multiple mating in *D. birchii* was associated with a reduction in the per mating number of offspring produced in later mating events (Fig. 1). This is also observed in *D. melanogaster* (Lefevre and Jonsson, 1962), despite substantially lower absolute productivity per mating in *D. birchii*. Importantly, these patterns were not an artefact of the non-random subset of males for which estimates of fourth matings are possible (i.e. those males that successfully copulate four times) because a comparable reduction in productivity across matings was seen for this subset of males too (Fig. SM1). This suggests rapid ejaculate depletion even in males that successfully mated four times, and contrasts with studies in tropical tephritid flies that found males do not deplete sperm stores, allocating similar quantities over three consecutive matings (Perez-Staples and Aluja, 2006).

A consistent increase in mean mating duration was observed with each successive copulation, from ∼3 to 6 minutes over four matings (Fig. 1). Extending mating durations can increase male fitness by increasing paternity (Bretman *et al*, 2013a; Mazzi *et al*, 2009). By contrast, in several other *Drosophila* species copulation durations decline over consecutive matings (Linklater *et al*, 2007; Singh and Singh, 2013; Singh and Singh, 2000), suggesting that duration decreases as male ejaculate becomes depleted (Edvardsson and Canal, 2006). In these experiments however, declining productivity with increasing duration indicates that extended copulation duration is not associated with increased sperm transfer. Gilchrist and Partridge (2000) found that sperm transfer occurs rapidly after the initiation of mating in *D. melanogaster* and that extended copulations delay female remating by transmitting seminal fluids that boost fecundity and reduce receptivity in females. Similarly, increases in mating duration with repeated male copulations in the housefly (*Musca domestica*), were likely to be due to the depletion of such accessory secretions, with males only terminating copulations when the levels transferred were sufficient to stimulate an inhibitory response to further mating in females (Leopold *et al*, 1971). While mating duration is largely controlled by males in several *Drosophila* (MacBean and Parsons, 1967; Parsons and Kaul, 1966; Patty, 1975; Singh and Singh, 2000), studies with *D. birchii* have shown that females can dislodge males (Hoikkala and Crossley, 2000). Similarly, females have an influence on copulation duration in *D. mojavensis* (Krebs, 1991), *D. elegans* (Hirai *et al*, 1999) and *D. montana* (Mazzi *et al*, 2009). Further research is necessary to clarify female fitness effects of prolonging copulations. Alternatively, prolonging copulation might represent a mate guarding strategy by males, particularly as sperm becomes limited (Simmons, 2001). However, the relatively small absolute increases in mean copulation duration in this study make such hypothesised mate guarding unlikely.

Males were assayed under standardised (constant 25 °C) laboratory conditions. However, male allocation strategies are likely to show adaptive responses to the more variable conditions experienced in space and time by natural populations (Wedell *et al*, 2002; Wigby *et al*, 2016). Pitnick and Markow (1994) propose that submaximal male ejaculate allocation over successive matings may act as bet-hedging for male *Drosophila*, particularly where environments are stressful. Variation in abiotic conditions also affects insect mating duration (Horton *et al*, 2002), sperm allocation and remating rate (Katsuki and Miyatake, 2009) within genotypes. Latitudinal variation in male remating traits within two cosmopolitan *Drosophila* species has also been observed (Chahal *et al*, 2013; Singh and Singh, 2000). Bouletreaumerle *et al* (1982) found that female fecundity was lower in tropical versus temperate populations of *D. melanogaster*, suggesting that males of tropical species may also invest less in sperm or seminal components associated with the environments they inhabit.

Despite this evidence for adaptive divergence in male trait in response to the local environment, we found no evidence for genetic divergence in any mating trait across either elevational gradient (Table 1), even though high and low elevation thermal environments (∼7 °C difference in mean temperature) characterise the cold and warm ecological limits of this species. However, there was significant genetic variation between isofemale lines in number of offspring (productivity) over successive matings at both gradients, with between line variance representing ∼17% of total variance in Mount Edith lines and ∼4% at Mount Lewis. Similar among-line variation was observed for mating duration at Mount Edith although not at Mount Lewis. Broad-sense heritability (*H*^2^) estimates varied substantially between gradients with Mount Edith showing consistently lower levels of genetic variance. For instance, offspring per mating gave values of 0.229 at Mount Edith compared to 0.069 at Mount Lewis. However, latency to mate showed no between-line variation and mating duration was only significantly heritable at Mount Edith.

Despite the restricted distribution of *D. birchii*, and the strong effects of these traits on fitness, heritabilities are remarkably high, at least at Mount Edith, compared to the wide-range of estimates obtained in *D. melanogaster*, for male courtship traits (0.033 – 0.094) (Gaertner *et al*, 2015), mating latency (0.01), duration (0.007) (Gromko, 1987) and mating success (0.25) (Tucic *et al*, 1988). Taylor *et al* (2013) observed no significant heritable variation in male mating latency and duration. Maintenance of genetic variation in such key fitness traits may depend on potential trade-offs between investment in sperm/seminal fluids and resistance to environmental stress. For example, thermal stress can have a substantial detrimental impact on male reproduction in ectotherms, including *D. birchii*, possibly due to the need to divert resources away from sperm production to upregulate physiological stress responses (Sales *et al*, 2018; Saxon *et al*, 2018).

Sexual conflict, ‘good genes’ and ‘sexy sons’ models of sexual selection all predict that males pass on genetic benefits to offspring (Kokko, 2001; Taylor *et al*, 2013; Wedell and Tregenza, 1999). However, if decreasing ejaculate quantity over successive matings in males is associated with declines in sperm quality, this may affect the fitness of offspring from later matings, making it easier to explain such high levels of variation in male mating strategies, even when they increase the number of first-generation offspring produced. However, Experiment II demonstrated that the paternal (sire) mating sequence had no significant effect on the reproductive success of sons from later matings compared to those from first mating, when they were mated to a single female (Fig. 4). This means that no deterioration in sperm/ejaculate quality is observed at least until after males’ fourth mating, which is associated with a 237% relative increase in mean total productivity at Mount Edith and 111% increase at Mount Lewis compared to a male that mates only once. Such variation in male remating rate within *D. birchii* populations therefore remains unexpected, given the apparent unequivocal fitness advantage it provides.

## Acknowledgements

We are grateful to Rhonda Snook for work assaying *D. birchii* sperm morphology and female reproductive tracts and to Tom Tregenza for helpful comments on a draft of the manuscript. This work was funded by a University of Bristol PhD scholarship for ADS and a Natural Environment Research Council standard grant (no. NE/G007039/1) to JRB.

## Data Archiving

All data referred to in this manuscript will be archived in Dryad.

