## Supplementary Material for "Low but genetically variable male remating ability in a tropical *Drosophila* despite substantial fitness benefits"

*(a) Formula for $H^2$ calculation*

$H^2$ was calculated following the method of Hoffmann and Parsons (1988) as:

$$H^2 = \frac{1}{2F_t} V_b / (V_b + V_w) \qquad (1)$$

where $F_t$ is the inbreeding coefficient after *t* generations as isofemale lines, calculated according to Falconer and Mackay (1996) as:

$$F_t = \frac{1}{2N} + \left(1 - \frac{1}{2N}\right) F_{t-1} \qquad (2)$$

where *N* is the population size in each generation. We calculated $F_t$ assuming a population size of 100 in each generation after establishment, and 25 generations at Mount Edith and 50 generations at Mount Lewis. We assumed that offspring in the first generation were all full-sibs (*i.e.* $N_{Gen0} = 2$; $F_0 = 0.25$).

*(b) Number of offspring produced by males achieving four matings*

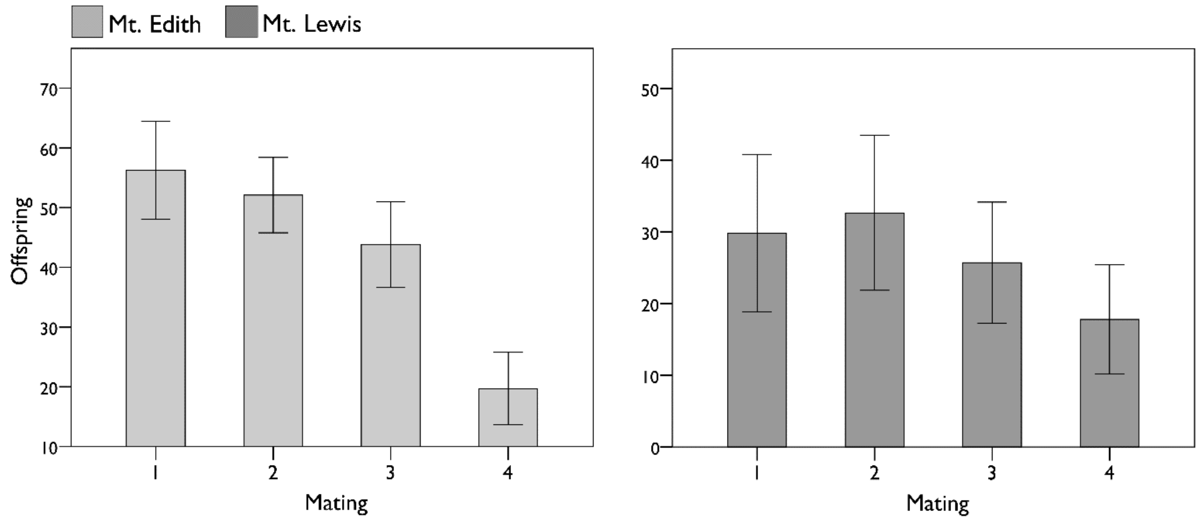


**Figure SM1. N**umber of offspring produced, calculated using by the subset of males reaching a fourth mating
